# Deep Learning Bridges Histology and Transcriptomics to Predict Molecular Subtypes and Outcomes in Muscle-Invasive Bladder Cancer

**DOI:** 10.1101/2025.10.23.684013

**Authors:** Alice Blondel, Clémentine Krucker, Valentin Harter, Melissa Da Silva, Clarice S. Groeneveld, Aurélien de Reynies, Maryam Karimi, Simone Benhamou, Isabelle Bernard-Pierrot, Christian Pfister, Stéphane Culine, Yves Allory, Thomas Walter, Jacqueline Fontugne

**Affiliations:** Centre for Computational Biology (CBIO), Mines Paris, PSL University, 75006, Paris, France; Institut Curie, PSL University, 75005, Paris, France; INSERM, U900, 75005, Paris, France; Institut Curie, PSL Research University, CNRS, UMR144, Equipe Labellisée Ligue Contre le Cancer, Paris, France; North-West Canceropole Data Center, Comprehensive Cancer Center Francois Baclesse, UNICANCER, Caen, France; Centre de Recherche des Cordeliers, Université Paris-Cité, UMRS1138, Paris; Bureau Biostatistique et Epidémiologie, Institut Gustave Roussy, Université Paris-Saclay, CESP U1018 Oncostat, Villejuif, France; Department of Urology, Charles Nicolle University Hospital, Rouen, France; Clinical Investigation Center, Onco-Urology, Inserm 1404, Rouen, France; Medical Oncology Department, Saint-Louis Hospital, AP-HP.Nord Université Paris Cité, Paris, France; Université Paris-Saclay, UVSQ, Institut Curie, Department of Pathology, Saint-Cloud, France

**Author notes:** All authors contributed equally (Thomas Walter, Jacqueline Fontugne). **Corresponding authors:** Alice Blondel, Thomas Walter, Jacqueline Fontugne.

**Keywords:** Deep Learning, Gene expression prediction, Molecular Subtyping, Intratumoral Heterogeneity, Muscle-Invasive Bladder Cancer (MIBC), Prognostic Implications, Spatial Transcriptomic

## Abstract

Muscle-Invasive Bladder Cancer (MIBC) is a heterogeneous disease with distinct molecular subtypes influencing prognosis and therapeutic response. However, molecular profiling through RNA sequencing remains costly, time-consuming and complicated by intratumoral heterogeneity. We developed a Deep Learning (DL) approach to infer molecular subtypes from routine histopathological slides and to evaluate its prognostic value in patients treated with neoadjuvant chemotherapy (NAC). We developed an DL-model predicting the expression of 848 subtype-associated genes from histological images of transurethral resection of bladder tumor, enabling spatial molecular subtyping at tile level. The model was trained on 297 NAC-treated patients from the VESPER clinical trial and evaluated on three independent cohorts (COBLAnCE, n=224; Saint-Louis, n=30 and TCGA, n=315), covering diverse staining protocols and scanner types. Spatial transcriptomics from six VESPER patients confirmed the spatial consistency of the inferred expression profiles. Our approach achieved a ROC AUC of 0.94 for molecular subtype prediction, with 95% of genes significantly predicted, demonstrating its ability to capture transcriptomic dysregulations from histological morphology. Predicted expression maps revealed spatially coherent patterns and intratumoral molecular heterogeneity. Importantly, tumors predicted with basal/squamous features (pure or mixed), were associated with significantly worse progression-free and overall survival after NAC (log-rank p=0.014 and 0.037, respectively). This DL-based framework enables accurate and spatially resolved inference of gene expression and molecular subtypes in MIBC without sequencing. These findings could improve patient stratification in clinical practice and support the design of more targeted clinical trials. Further validation in larger cohorts is needed before routine clinical implementation.

## 1. Introduction

Muscle-invasive bladder cancer (MIBC) is an aggressive and heterogeneous disease, accounting for approximately 25% of newly diagnosed bladder cancers [1]. It is associated with a high rate of recurrence and mortality, with a 5-year survival rate of 40-60% [2,3]. The current therapeutic standard - neoadjuvant chemotherapy (NAC) followed by cystectomy - offers limited benefit for many patients, partly due to the biological diversity of the disease.

Transcriptomic studies have revealed that MIBC encompasses multiple molecular subtypes, such as luminal, basal-squamous (Ba/Sq), and neuronal [4–6]. To unify classification systems, a Consensus Molecular Classification was proposed [7], providing a robust framework to stratify MIBC into clinically relevant subgroups. Growing evidence suggests that these subtypes are associated with distinct clinical outcomes and differential treatment responses [8,9]. In particular, the Ba/Sq subtype is frequently linked to tumor aggressiveness and poor prognosis [10,11].

While bulk RNA sequencing remains the gold standard for molecular subtyping, its clinical implementation is limited due to cost, turnaround time, and infrastructural demands. Moreover, many MIBCs exhibit intratumoral molecular heterogeneity, with multiple subtypes coexisting within the same tumor [12], contributing to poor NAC response and worse survival [10]. These challenges highlight the need for scalable, cost-effective, and spatially resolved tools, for molecular stratification from routine clinical specimens.

Advances in Deep Learning (DL) applied to histopathology have opened avenues for inferring molecular information directly from digitized hematoxylin-eosin (HE)-stained slides [13,14]. In MIBC, studies have demonstrated the feasibility of predicting molecular subtypes from whole-slide images (WSIs) [15–17]. While promising, these approaches often suffer from limited spatial resolution, coarse subtype granularity, or reliance on traditional machine learning models, and survival analyses rarely assess the prognostic value of histology-inferred subtypes.

Recent studies in digital pathology have demonstrated that predicting gene expression from histology enables more accurate and interpretable models, supporting diverse downstream biological and clinical applications [18–20].

Here, we introduce a state-of-the-art DL-based model predicting the expression of 848 genes associated with the Consensus MIBC Classification [7] from histological images. By predicting gene expression at the tile level - tiles correspond to patches of 112 µm - our method enables fine-grained spatial mapping of molecular subtypes, thereby uncovering intratumoral heterogeneity. We evaluated this approach across three independent MIBC cohorts and used spatial transcriptomic (ST) data to validate the spatial consistency and biological relevance of the inferred expression profiles (Figure 1A).

**Figure 1:**
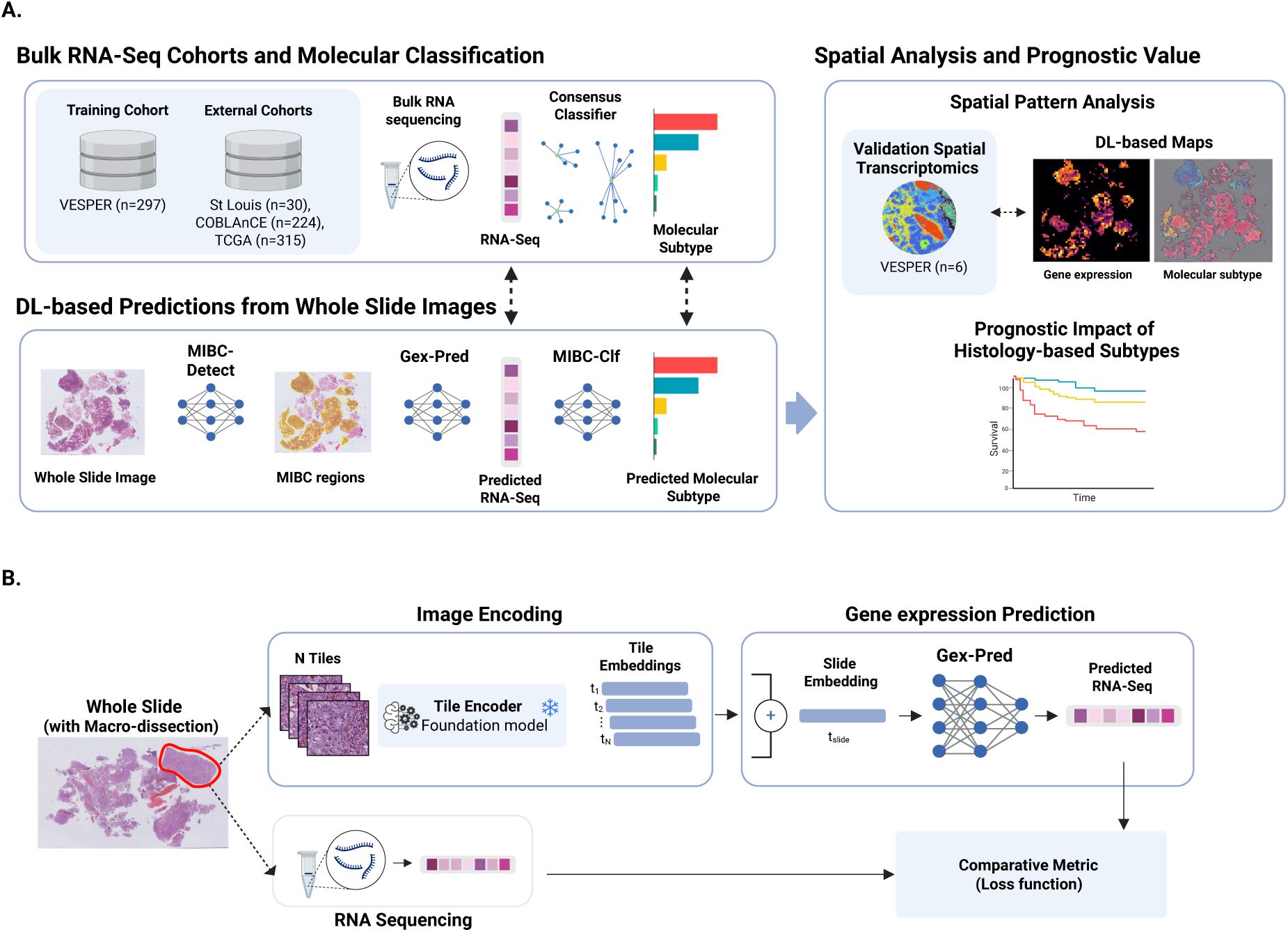
Methodology for molecular subtype prediction in MIBC. **A.** Overview of the proposed AI-based pipeline. It consists of three models: MIBC-Detect to identify MIBC tumor regions, GeX-Pred to predict gene expression from those regions, and MIBC-Clf to classify molecular subtypes. The pipeline is trained and validated using matched whole-slide images (WSIs) and bulk RNA-seq data from the VESPER cohort. External validation is performed on three independent cohorts (Saint Louis, COBLANCE, and TCGA). In addition to tissue-level gene expression prediction, spatial predictions are integrated and validated using spatial transcriptomics. The clinical relevance is demonstrated through survival analysis based on molecular subtype stratification. **B.** Architecture and training pipeline of GeX-Pred. Tissue sections are divided into tiles and encoded using a foundation model. GeX-Pred is trained so that the mean expression predicted across tiles matches the bulk RNA-seq profile.

## 2. Materials and Methods

### 2.1. Description of the cohorts

This study leveraged one training cohort (n=297) and three independent validation cohorts (n=569), collected across mono- and multicentric settings. The cohorts cover diverse clinical protocols, tissue processing methods, and scanner types, allowing evaluation across heterogeneous real-world conditions. A summary of key cohort characteristics is provided in Supplementary Figure 1.

The training dataset was derived from the VESPER cohort (NCT01812369), a multicentric prospective phase III clinical trial [21] enrolling cT2N0 MIBC patients, treated with NAC. We selected 297 patients for which formalin-fixed paraffin-embedded (FFPE) hematoxylin-eosin-safran (HES)–stained slides from transurethral resection of bladder tumor (TURBT) specimens and matched RNA-seq were available.

For validation, three external cohorts were used:(i) a retrospective monocentric cohort from Saint-Louis Hospital, comprising 30 MIBC patients treated with NAC; (ii) 224 patients from the COBLAnCE prospective multicentric cohort [22], including only MIBC cases (both NAC-treated and NAC-naive) originating from 13 institutions, with FFPE histological slides from TURBT and/or cystectomy specimens; and (iii) 315 MIBC patients from The Cancer Genome Atlas (TCGA) [23], with FFPE TURBT or cystectomy slides reviewed by a pathologist to ensure inclusion criteria were met (see Supplementary Section 1.b).

For all cohorts, inclusion required urothelial MIBC, FFPE slides, and molecular data passing quality control. Tumors with neuroendocrine differentiation or classified as “NE-like” by molecular profiling, were excluded, as NAC is not standard of care for this subtype.

### 2.2. Transcriptome profiling and molecular subtyping

Transcriptomic profiling was performed on FFPE or frozen samples. In VESPER, Saint-Louis, and COBLAnCE FFPE cohorts, homogeneous tumor areas were macro-dissected - to enrich for cancer cells and ensure homogeneity - and sequenced using 3ʹ-seq or SMARTer protocols. RNA-Seq data were obtained from frozen samples in COBLAnCE Frozen and TCGA datasets. Data were normalized and consensus molecular subtypes were assigned using the “consensusMIBC” R package [7]. ST was generated for six VESPER patients using the Visium HD protocol. Detailed cohort-specific information is provided in Supplementary Section 1.b and Supplementary Figure 1.

### 2.3. Slide scanning

All VESPER, COBLAnCE FFPE and Saint-Louis blocks were recut and stained at Curie Institute using HES staining. VESPER comprised 514 HES-stained slides. COBLAnCE FFPE subset and Saint-Louis cohorts comprised 80 and 30 HES-stained slides, respectively. All VESPER, COBLAnCE, Saint-Louis and Visium slides were digitized at 40X magnification using Hamamatsu NanoZoomer S360MD scanners at Curie Institute (see Supplementary Figure 1). COBLAnCE Frozen cohort included 147 patients, with FFPE slides originally stained in HES or HE; the set comprises both original and re-stained slides scanned at Curie, totaling 597 slides. Publicly available TCGA slides were scanned across multiple institutions, and all were HE-stained [6].

### 2.4. DL-Based Gene Expression and Molecular Subtype Prediction

We developed a multi-stage DL framework to predict molecular subtypes from WSIs. The framework comprises three components: Gex-Pred (gene expression predictor), MIBC-Clf (molecular subtype classifier), and MIBC-Detect (tumor region selector; Figure 1A,B). The framework leverages recent advances in computational pathology, including transformer-based feature extraction [24,25], transfer learning [26], and weakly supervised learning [27,28] (see Supplementary Section 2).

WSIs were divided into non-overlapping 224×224-pixels tiles at 20X magnification (∼ 112 μm) and encoded using H-optimus-1 - a model pretrained on millions of histology slides - to extract rich and transferable features.

Gex-Pred inferred the expression of the 848 subtype-associated genes. Using an attention-based multiple instance learning (MIL) strategy [27], Gex-Pred aggregated tile-level representations into a slide-level representation, which was then used to predict bulk RNA expression (Figure 1B and Supplementary Section 2). Based on the predicted expression profiles, MIBC-Clf assigned consensus molecular subtypes.

MIBC-Detect was trained to distinguish MIBC tiles from non-MIBC and non-tumor tiles using manual annotations from the VESPER cohort provided by a senior pathologist with QuPath [29] (see Supplementary Section 2.c). Subtype predictions were then restricted to regions identified as MIBC by MIBC-Detect.

### 2.5. Performance assessment and statistical methods

Model evaluation on the VESPER cohort was performed using 10-fold patient-wise cross-validation. The accuracy of Gex-Pred for transcriptomic inference was measured by the Pearson correlation coefficient (PCC) between predicted and bulk RNA-seq expression profiles. Statistical significance was assessed using two-sided tests with a significance threshold of 0.05, adjusted for multiple comparisons via the Benjamini-Hochberg method. The performance of MIBC-Clf for molecular subtype prediction, as well as MIBC-Detect for MIBC region identification, was evaluated using the area under the receiver operating characteristic curve (ROC-AUC), with standard deviations reported across cross-validation folds.

Survival differences between molecular subgroups were analyzed using Kaplan–Meier survival curves and multivariate log-rank tests to determine statistical significance.

Finally, the trained model was frozen and applied to external validation cohorts (Saint Louis, COBLAnCE, TCGA, and ST) to assess its generalizability.

## 3. Results

### 3.1. Accurate prediction of Gene expression and Molecular subtypes

Performance of Gex-Pred, predicting 848 subtype-associated genes, is summarized in Figure 2 across cohorts. In macro-dissected VESPER samples, the median PCC reached 0.46, with 95% of genes significantly predicted. Comparable performance was observed in external macro-dissected cohorts (COBLAnCE: median PCC 0.57; Saint Louis: median PCC 0.60, with 25% of genes exceeding 0.70). Performance remained robust on frozen samples (median PCC of 0.46 and 0.45 for COBLAnCE and TCGA, respectively).

**Figure 2:**
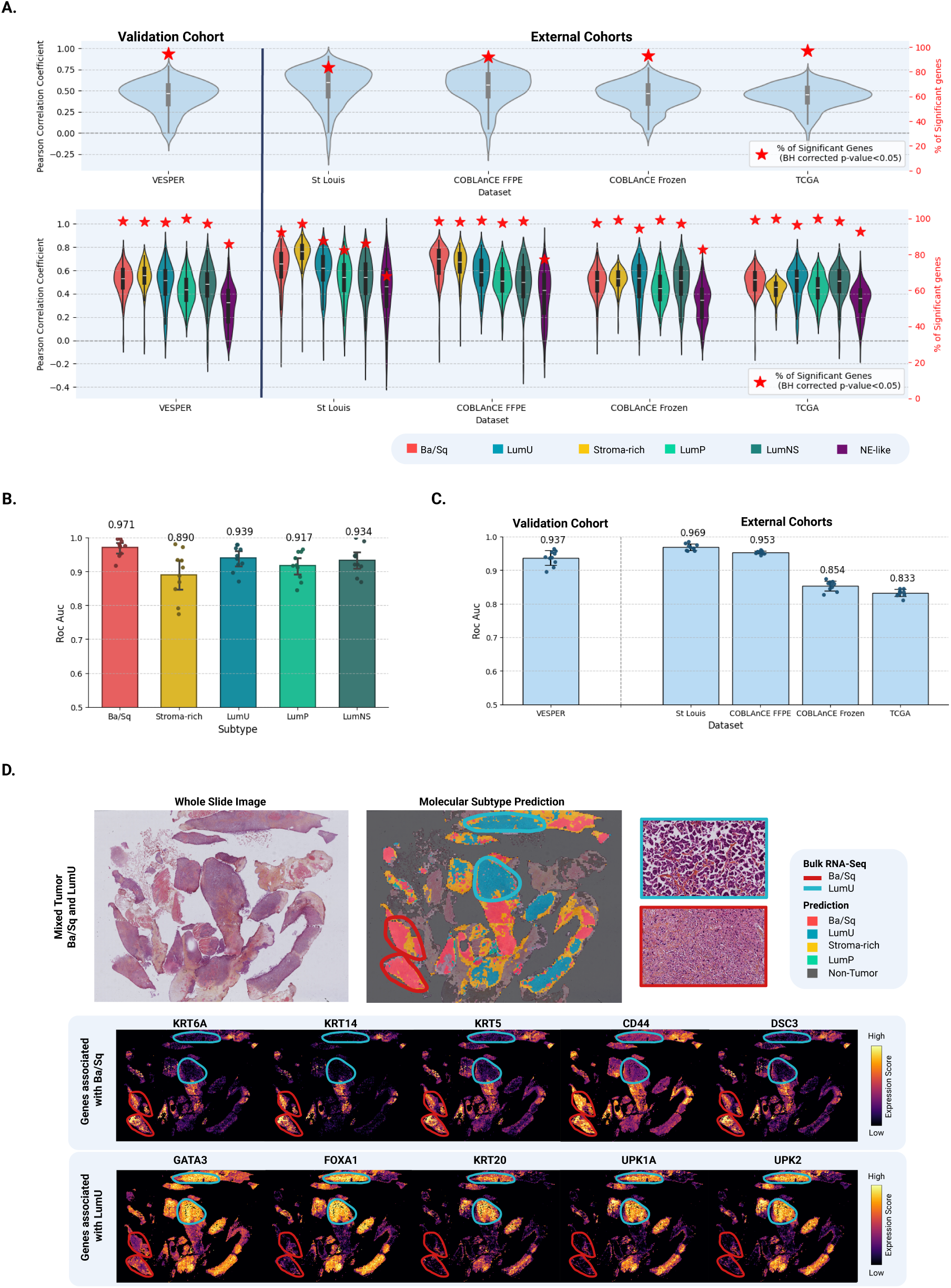
Performance assessment of gene expression and molecular subtype prediction. **A.** Gene expression prediction performance for the 848 consensus MIBC genes across cohorts. Violin plots show the distribution of Pearson correlation coefficients (left y-axis) between predicted and actual gene expression. Red stars indicate the percentage of genes (right y-axis) with significantly well-predicted expression. A second row displays the same metrics, grouping genes by molecular subtype association, to assess prediction performance across biological classes. **B.** Molecular subtype classification performance on the VESPER cohort. ROC-AUC and confidence intervals are reported for each subtype. **C.** Molecular subtype classification performance across external cohorts (Saint Louis, COBLAnCE, and TCGA). ROC-AUC values are reported per cohort. **D.** Spatial visualization of predicted gene expression and molecular subtypes for a mixed tumor of VESPER cohort. The spatial maps reveal co-occurrence of LumU and Ba.Sq subtypes with distinct morphologies. Tile-level predictions are shown for five known LumU marker genes and five Ba.Sq marker genes.

Ba/Sq-associated genes showed the highest correlations, reflecting subtype-specific morphological-transcriptomic patterns (see Figure 2A), whereas neuroendocrine (NE) genes were poorly predicted, consistent with NE cases being excluded from training (Supplementary Section 2.f for gene-subtype assignment).

MIBC-Clf accurately predicted consensus molecular subtypes, achieving an overall ROC AUC of 0.937 on VESPER samples. Ba/Sq subtype preformed best (ROC AUC of 0.971), with all subtypes above 0.890 (Figure 2B,C). Independent macro-dissected cohorts (Saint-Louis and COBLAnCE) maintained strong performance (ROC AUC of 0.969 and 0.953, respectively), demonstrating good generalizability across centers (Figure 2C). Performance slightly dropped in cohorts with RNA-seq from adjacent frozen samples (0.854 and 0.833 for COBLAnCE Frozen, and TCGA respectively), reflecting spatial mismatch and intratumoral heterogeneity. In COBLAnCE Frozen, model performance was compared on TURBT and cystectomy specimens, with a ROC AUC of 0.876 for cystectomy confirming its applicability to this tissue type (Supplementary Figure 4).

MIBC-Detect achieved a ROC AUC of 0.978 (Supplementary Figure 6) in VESPER cohort, enabling precise MIBC localization within WSIs before molecular subtype classification.

Thus, we demonstrated that our model faithfully reproduces bulk-level gene expression predictions and associated molecular classifications. Applied at the tile level across whole slides, it generates spatially resolved prediction maps illustrating the intratumoral distribution of molecular subtypes (Figure 2D).

### 3.2 ST validates the spatial predictions

To validate the model’s spatial prediction performance at the tile level, we used ST data generated with Visium HD technology. Tiles of 112 µm were extracted via a sliding window (stride: 100 µm), and both Gex-Pred and MIBC-Clf were applied to each tile. Predictions were compared to ground truth derived by aggregating ST gene expression within the corresponding regions. As shown in Figure 3D,E, the model accurately identified tissue subtypes and gene expression across samples. We observed a median PCC of 0.233 for gene expression, reflecting the technological gap between bulk RNA-seq (used for training) and ST (used for evaluation). Despite this, the model achieved robust molecular prediction, with a ROC AUC of 0.870. For three patients (Figure 3A-C), we visualized the spatial expression of two subtype-specific genes. The model accurately recapitulated gene spatial distribution. Additional examples across more slides and genes are presented in Supplementary Figure 5.

**Figure 3:**
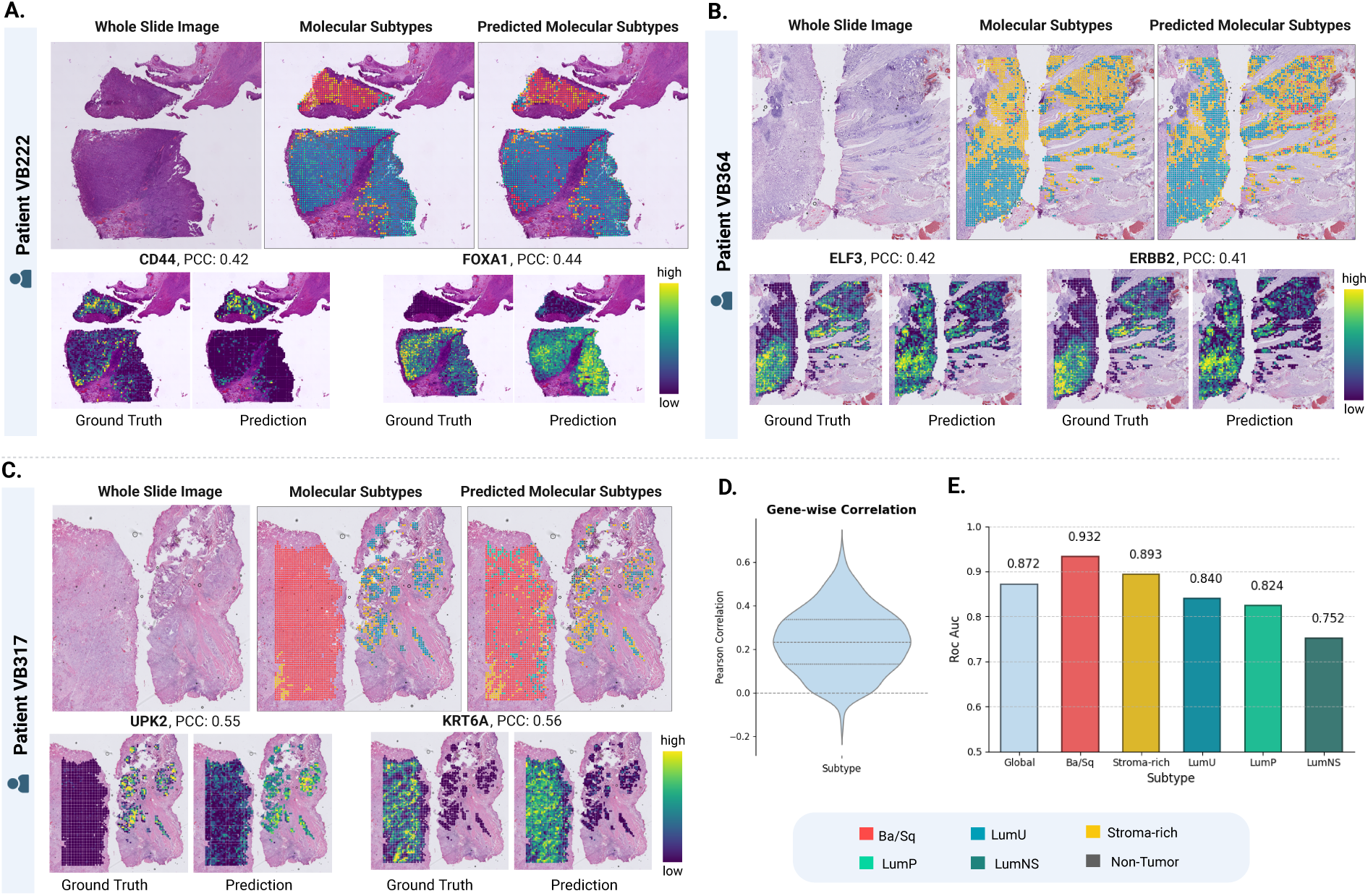
Validation of gene expression and molecular subtype predictions using ST. **A–C:** Comparison of model predictions with ground truth for three patients from the VESPER cohort. For each patient, the first row shows: (i) the WSI thumbnail, (ii) molecular subtype distribution derived from ST, and (iii) subtype classification predicted by MIBC-Clf. Two subtype-specific genes were selected per patient (ST data and the corresponding prediction by GeX-Pred). The PCC between predicted and actual spatial expression is indicated. **D:** Distribution of gene-wise PCC across all ST samples. **E:** ROC AUC for molecular subtype classification, showing both global AUC and per-class AUC across all ST samples.

These results demonstrate that histomorphology encodes rich molecular information, enabling reliable, spatially resolved prediction of tumor subtypes across diverse technologies.

### 3.3 Interpretability and Robustness of the model

Our approach can predict gene expression at the tile level, making it interpretable by design. It identifies specific regions on WSIs that display phenotypes associated with each molecular subtype and captures the intratumoral spatial organization, thus revealing spatial heterogeneity (Figure 2D). Gene-level prediction maps reveal the spatial distribution of key molecular markers, offering direct insights into the transcriptomic features underlying subtype assignments.

Additionally, interpretability was enhanced by examining tiles with the highest prediction scores for each molecular subtype (Supplementary Figure 7A). These tiles often exhibited morphological features consistent with known subtype-associated phenotypes, supporting the biological relevance of the model’s predictions.

The model’s robustness was demonstrated by its consistent performance on consecutive slides from the COBLAnCE cohort (Supplementary Figure 7B) processed at different centers, highlighting its resilience to staining variability. Importantly, we observed a strong correlation (Pearson r = 0.954) between the subtype tile compositions predicted on matched slides from different centers (n=142). Robustness across technical variations is an essential prerequisite for future clinical translation.

### 3.4 DL highlights intratumor heterogeneity and its prognostic impact

Using our DL model, we accurately classified macro-dissected samples from the VESPER cohort into the five molecular subtypes. We further investigated spatial patterns of molecular subtype and tumor heterogeneity (Figure 4A), identifying both homogeneous tumors and “mixed” tumors composed of multiple subtypes based on macro-dissected tumor regions.

**Figure 4:**
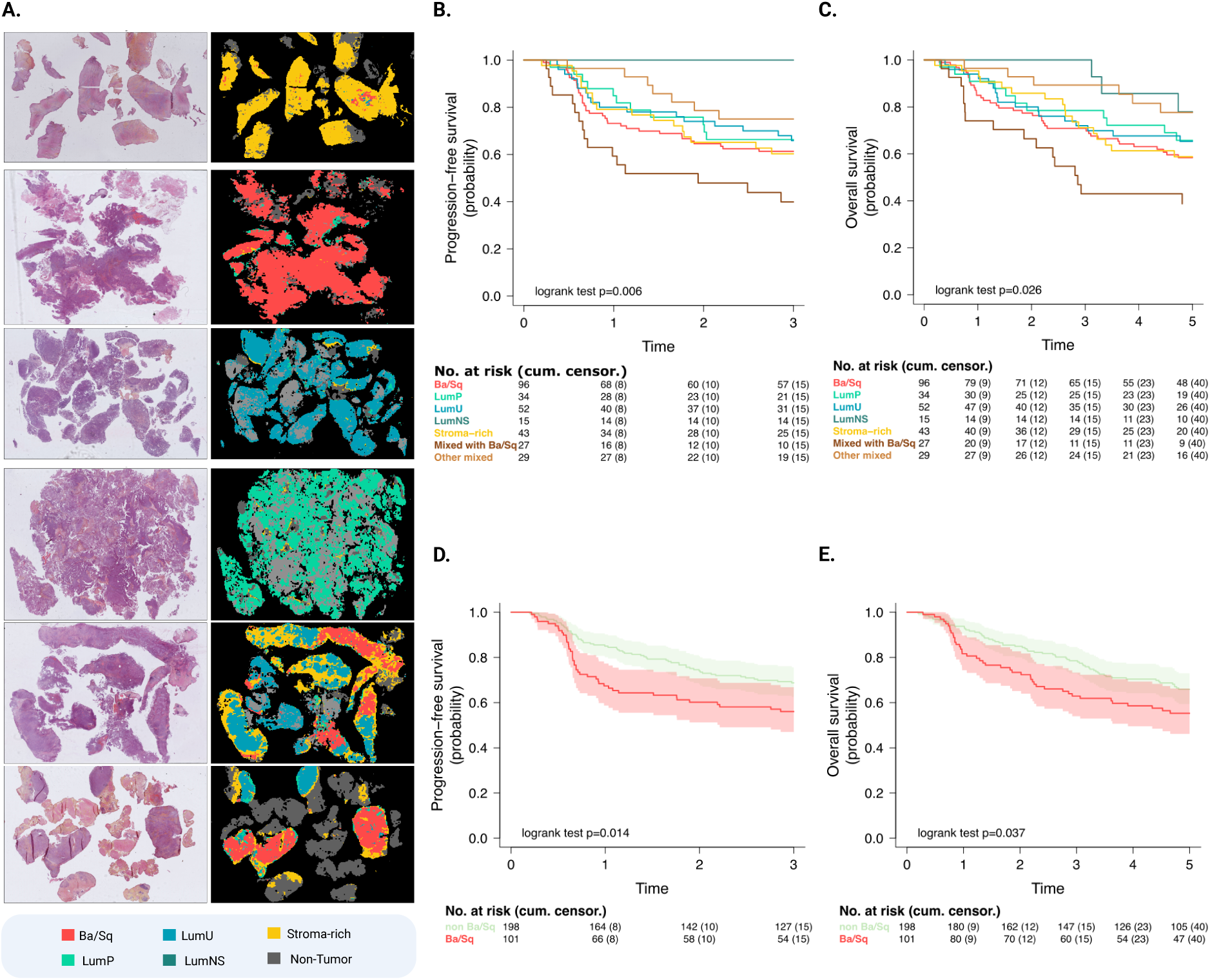
Prognostic impact of AI-based molecular stratification and intratumoral heterogeneity. **A)** Representative examples of homogeneous tumors (single subtype) and mixed tumors (multiple subtypes) highlighting spatial heterogeneity. **B–C)** Kaplan–Meier survival curves for progression-free survival (B) and overall survival (C) based on predicted stratification into five Consensus MIBC subtypes and mixed tumors. **D–E)** Kaplan–Meier survival curves for whole-slide patient stratification into Ba/Sq versus non-Ba/Sq tumors (D, PFS; E, OS).

To evaluate the prognostic impact of this heterogeneity, patients were stratified into seven groups: five homogeneous subtypes, a “Mixed” group (multiple subtypes without Ba/Sq), and a “Mixed + Ba/Sq” group (mixed tumors containing a Ba/Sq component). Overall, 19% of patients exhibited subtype heterogeneity, including 9% in the Mixed + Ba/Sq category. Kaplan-Meier survival analysis revealed that tumors with a basal component had significantly worse outcomes than all other groups (log-rank PFS = 0.006, OS = 0.026) (Figure 4B,C). Additional analyses stratifying tumors into two groups - Ba/Sq vs non-Ba/Sq - yielded consistent prognostic trends (Supplementary Figure 8). These results highlight that spatial intratumoral heterogeneity carries strong prognostic significance in the NAC setting.

### 3.5 A clinically relevant tool

To evaluate the clinical applicability of our approach, we applied Gex-Pred to all available whole-slide images, generating patient-level gene expression predictions across entire tumors. We subsequently applied MIBC-Clf to stratify patients into two groups: Ba/Sq and non-Ba/Sq. This simplified classification retained strong prognostic value, with Kaplan-Meier analyses demonstrating significant survival differences between the two groups (log-rank PFS = 0.014, OS = 0.037) (Figure 4). These results indicate that our model could serve as a clinically actionable tool, enabling robust risk stratification from standard histology alone and potentially guiding therapeutic decisions in the NAC setting.

## 4. Discussion

NAC remains the standard of care for MIBC, yet response prediction remains a major clinical challenge. In this study, we demonstrate that molecular subtypes and gene expression profiles can be accurately inferred from standard histology slides using an interpretable DL framework. By leveraging routine H&E slides, our model provides a cost-effective and scalable alternative to molecular profiling, potentially enabling molecular-based patient stratification in real-world settings.

Our findings confirm that the morphological patterns visible in MIBC histology encode robust molecular information, consistent with previous reports linking histopathology and transcriptomic features [15,17]. Trained on macro-dissected regions spatially matched to RNA sequencing, our approach integrates precise supervision with large-scale multi-institutional data, resulting in high generalizability across scanners, stains, and cohorts. The model reproduced consensus molecular subtype distributions across VESPER, COBLAnCE, Saint-Louis and TCGA cohorts, and showed spatial coherence with Visium HD data, suggesting that it captures biologically meaningful intratumoral heterogeneity.

Importantly, the distribution of DL-predicted subtypes carried significant prognostic implications. Ba/Sq tumors, as predicted from histology, were associated with poor outcome after NAC, consistent with prior molecular studies [7,10,11]. Moreover, mixed tumors containing Ba/Sq areas displayed even worse prognosis, underscoring the clinical relevance of heterogeneity analysis. To our knowledge, this is the first work directly assessing the prognostic impact of DL-predicted subtypes in MIBC, while most previous studies focused on agreement between DL predictions and molecular labels [15,17].

From a methodological standpoint, our approach moves beyond conventional subtype classification by first regressing full gene expression profiles and subsequently inferring molecular subtype. This formulation leverages the continuous nature of transcriptomic data, enriching the training signal, mitigating class imbalance, and enabling multi-task learning across hundreds of genes (n=848), as recently advocated in other cancers [18,19]. Consequently, the model jointly learns transcriptomic and subtype-related patterns without relying on predefined thresholds. This framework demonstrates the feasibility of virtual transcriptomics from histology, supporting a broad range of downstream molecular analyses.

This study has several limitations. First, survival analyses were performed in VESPER cohort. Independent validation in larger real-world cohorts will be necessary to confirm the generalizability of the prognostic value of DL-predicted subtypes and to strengthen clinical translation. Second, the definition of heterogeneity relied on pathologist-guided annotations from macro-dissection and IHC, which may limit scalability. Automated or semi-automated annotation methods could facilitate future implementation. Third, while our framework focuses on predicting molecular features, future developments should aim to identify patients unlikely to respond to NAC, enabling early therapeutic redirection.

Finally, extending the framework to integrate multimodal data - such as clinical variables, medical imaging, or genomic alterations - could further enhance model performance and clinical relevance by capturing complementary aspects of tumor biology and patient context, as demonstrated in [30]. In future developments, maintaining histology as the primary input will be essential to ensure scalability, and seamless integration into clinical workflows, ultimately supporting actionable and widely deployable predictions in precision oncology.

## 5. Conclusions

In summary, this study demonstrates the feasibility of predicting molecular subtypes and transcriptomic profiles directly from histology in MIBC. By bridging pathology and molecular profiling, this DL-based approach provides a clinically applicable tool for molecular risk stratification and personalized treatment planning, paving the way for improved patient management and more targeted clinical trials in bladder cancer.

## Declaration of Generative AI and AI-assisted technologies in the writing process

During the preparation of this work the authors used ChatGPT to improve the English language quality. After using this tool, the authors reviewed and edited the content as needed and take full responsibility for the content of the publication.

## Acknowledgements

We would like to thank Julia Thierry Francois, the COBLAnCE investigators, the VESPER trial investigators. We thank PRAIRIE for funding Alice Blondel’s PhD and the Translational Research Program in Oncology (INCa PRT-K) for their support.

## Data and Code Availability

Source code, pretrained weights, and example scripts for applying the model will be released on GitHub upon manuscript acceptance.

## Supplementary Materials

### 1. Dataset Construction

#### a. Description of the cohorts

This study leveraged one training cohort (VESPER) and three independent validation cohorts (Saint-Louis, COBLAnCE, and TCGA-BLCA), covering diverse clinical settings, tissue processing methods, and sequencing protocols. Key characteristics of all cohorts, including sample size, staining methods, sequencing protocols and molecular subtype distribution, are summarized in Supplementary Figure 1.

**Supplementary Figure 1:**
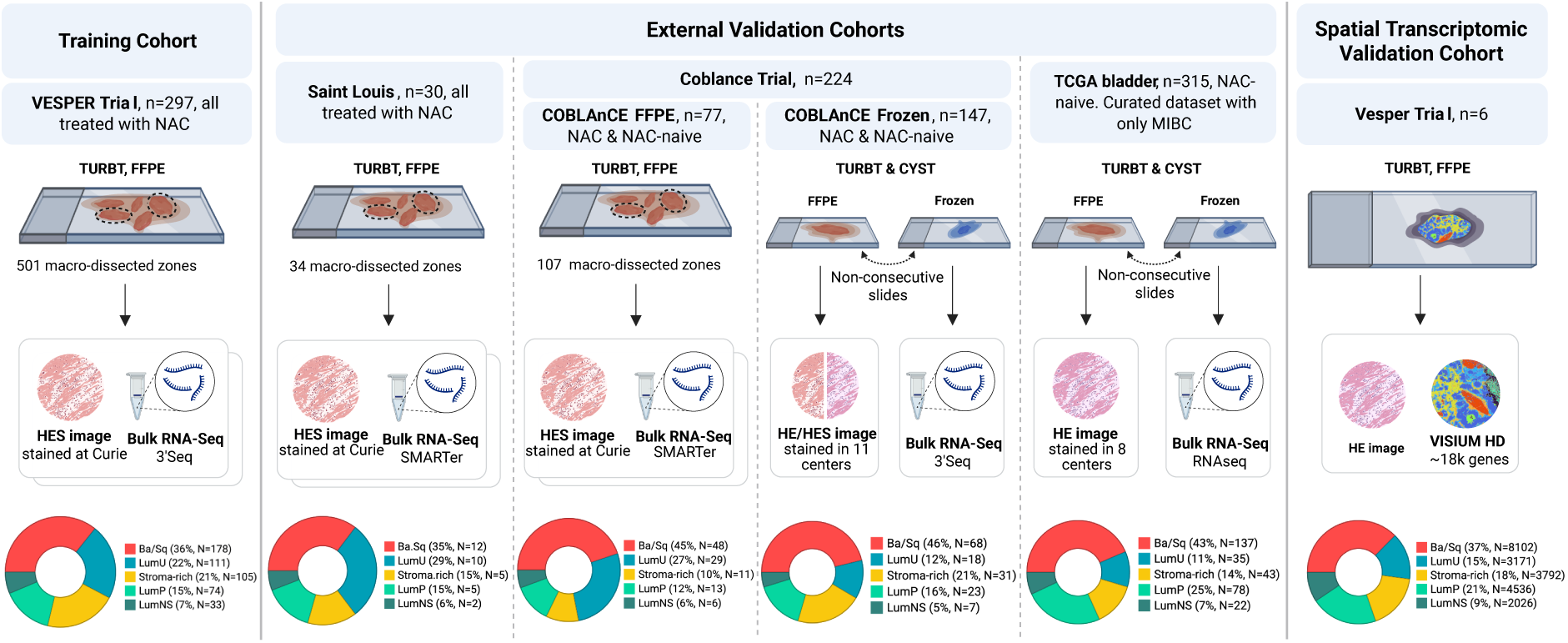
Overview of training and external cohorts. The training cohort consisted of 297 MIBC patients from the VESPER phase III trial, all treated with NAC, with HES-stained FFPE slides and bulk RNA-seq (3ʹSeq). A total of 501 tumor regions in VESPER were selected for macrodissection based on histological and immunohistochemical guidance. Validation cohorts included: (i) a monocentric series from Saint-Louis Hospital (n = 30, all treated with NAC, 34 regions sequenced using SMARTer); (ii) the prospective multicentric COBLAnCE study (n = 224 MIBC patients treated with NAC or NAC-naïve). The COBLAnCE FFPE subset (n = 77) includes HES-stained slides with RNA-seq performed using SMARTer protocols, while for COBLAnCE Frozen, 147 samples were sequenced using 3’Seq RNA-seq and (iii) the TCGA-BLCA cohort (n = 315), curated to include only MIBC cases, with HE-stained FFPE slides and frozen material for bulk RNA-seq. In addition, spatial transcriptomic validation was performed on six VESPER patients using the Visium HD platform (19,000 genes, 6.5 × 6.5 mm sections).

#### b. Transcriptome profiling and molecular subtyping

In the VESPER trial-derived cohort, bulk RNA-seq was performed on 415 macro-dissected tumor regions from 296 patients (GEO number: GSE279390), as previously described [10]. An additional 86 regions were macro-dissected using the same procedure to expand the training set for the deep learning algorithm, resulting in a total of 501 regions from 297 patients.

Briefly, to ensure cancer cell enrichment, homogeneous tumor areas were macro-dissected for RNA-sequencing and up to four regions were selected per patient in cases with morphological and/or immunohistochemical intratumoral heterogeneity. This macro-dissection strategy aimed to select spatially localized and molecularly homogeneous tumor regions. RNA was extracted and sequenced using the 3ʹ-seq protocol (QuantSeq 3’, Lexogen) and matched to corresponding histological images.

The Saint-Louis and COBLAnCE FFPE cohorts followed the same macrodissection-based methodology, using histological and immunohistochemical guidance. In the Saint-Louis cohort, 34 regions from 30 patients were sequenced using the SMARTer protocol. In the COBLAnCE cohort, 107 regions from 77 patients were sequenced using SMARTer protocols.

Additional transcriptomic data were available from frozen blocks in COBLAnCE and TCGA. In both cases, histological slides used for digital analysis were derived from FFPE tissue, which may not originate from the same tumor area as the one used for RNA extraction. In the COBLAnCE cohort, RNA profiling was performed using the 3ʹ-seq protocol, while the TCGA cohort relied on standard RNAseq.

Transcriptomic data were normalized using a log2 transformation of read-per-million (RPM) values, except for SMARTer-derived data, which were normalized using transcript-per-million (TPM) values. Consensus Molecular subtypes were subsequently assigned using the “consensusMIBC” R package [7]. The subtype distributions across cohorts are summarized in Supplementary Figure 1.

Finally, spatial transcriptomics (ST) data were generated for six VESPER patients using the Visium HD protocol, covering tissue sections of 6.5 × 6.5 mm. Gene expression data were aggregated over 100 μm-wide regions and log2-transformed. Consensus molecular subtypes were then assigned based on the normalized expression data.

#### c. Slide annotation for TCGA-BLCA cohort

This study focused specifically on MIBC, the tumor type underlying the development of the consensus molecular classification. Accordingly, the application of both the consensus classifier and Gex-Pred was restricted to MIBC cases. Given the potential discrepancies in tumor stage between frozen samples and matched adjacent FFPE slides, all available FFPE HE-stained slides from TCGA-BLCA cohort (n=386) were systematically reviewed by an expert genito-urinary pathologist (JF) (https://cancer.digitalslidearchive.org). Cases with FFPE slides for which the presence of MIBC could not be confirmed were excluded (n = 59). Specifically, we excluded 8 cases for which the FFPE slides did not harbor urothelial carcinoma and 51 cases which corresponded exclusively to an NMIBC component. This left 327 patients, of whom 4 lacked bulk RNA-seq data, 4 corresponded to the NE-like subtype, and 4 had only bulk RNA-seq data for Solid Tissue Normal samples (TCGA sample type 11A), resulting in a final cohort of 315 patients.

All available FFPE diagnostic slides associated with each case were included in the prediction workflow to capture spatial heterogeneity within the tumor and maximize representativeness of the molecular subtype assignment.

### 2. AI Methodology for spatial molecular prediction of MIBC

#### a. Preprocessing of whole-slide images: Segmenting, Tiling and Color Normalization

WSIs were acquired in SVS or NDPI format and uniformly downsampled to ×20 magnification. The application of deep learning to histopathological WSIs presents major computational and methodological challenges, primarily due to the extremely high resolution of the data (up to 100,000 × 100,000 pixels per slide, representing several gigabytes). To enable efficient learning and reduce dimensionality, we implemented a multi-step preprocessing pipeline.

We first identified tissue-containing regions within each WSI by applying Otsu thresholding on a down-sampled thumbnail, generating a binary tissue mask. Slides were then divided into smaller patches (tiles) of 112 × 112 μm (224 × 224 pixels at 0.5 μm/pixel resolution). Tiles were retained only if at least 50% of their area was classified as foreground (i.e., tissue) in the previous step.

To reduce staining variability, we applied color deconvolution [31] to convert HES-stained tiles into HE-like appearance by removing the saffron (S) component. This harmonization step improves the model’s robustness across cohorts by aligning staining protocols with the widely adopted HE standard. Tiles were then encoded into reduced-dimensional feature vectors using a pre-trained neural network to capture meaningful morphological information.

#### b. Tile Encoding with a foundation model

Due to the limited size of our dataset (501 macro-dissected regions from the VESPER cohort), training a deep neural network from scratch would be inefficient and prone to overfitting. Instead, we adopted a transfer learning approach inspired by recent advances in computer vision [32,33] and digital pathology[24,34–36].

Transfer learning leverages knowledge gained from a large source domain to improve performance on a specific target task. In our case, we used foundation models, large neural networks pre-trained on millions of histology tiles from diverse organs, staining protocols, and pathologies. These models, such as CTransPath [24], Phikon [34], UNI [35], Giga-Path [36] or H-optimus [37], have demonstrated strong generalization capabilities by capturing rich, task-agnostic representations of tissue architecture, including nuclear morphology, glandular structures, and spatial organization.

Specifically, we used H-optimus-1 as a feature extractor to encode each tile into a 1536-dimensional vector. H-optimus-1 is transformer-based architecture (ViTg/14, 1.1B parameters) trained in a self-supervised manner (Dinov2) on a million slides and 800 000 patients from various datasets, with extensive data augmentation (random cropping, flips, color jitter, grayscale, and Gaussian blur) and without any labels.

The feature extractor weights were frozen during both training and inference. After this step, each slide is represented as a matrix of size (number of tiles × 1536), which serves as input for subsequent prediction models.

#### c. MIBC-Detect Training

MIBC-Detect is a tile-level classifier that identifies tumor regions corresponding to MIBC. It distinguishes between three classes: MIBC tumor, non-MIBC tumor, and Other (including non-tumoral tissue, artifacts, pen marks, blurred regions). The model was trained on non-exhaustive annotations provided by expert pathologists on slides from VESPER cohort, comprising a total of 79,296 non-tumor tiles and 53,471 NMIBC tiles (see Supplementary Table 1). Tumor annotations correspond to the VESPER macro-dissected regions, along with 14 additional zones selected for their ambiguous phenotype, which is close to NMIBC but represents MIBC. Annotations were performed using QuPath.

**Supplementary Table 1:** Summary of tile-level annotations used to train the MIBC-Detect classifier. Each annotated region is assigned to one of three main classes: *MIBC tumor*, *Non-MIBC tumor*, or *Other*. The *Other* class includes a variety of non-tumoral tissues and structures (e.g., immune cells, muscle, connective tissue, fat), as well as artifacts and ambiguous regions. The table reports the number of annotated slides and the total number of extracted tiles per annotation type.

| Annotation | Class | # Slides | # Tiles |
| --- | --- | --- | --- |
| MIBC tumor | MIBC tumor | 435 | 874912 |
| NMIBC tumor | NMIBC | 60 | 53471 |
| Non-Tumor | Other | 18 | 51916 |
| Immune Cells | Other | 37 | 2038 |
| Urothelium | Other | 26 | 1930 |
| Muscle | Other | 25 | 6643 |
| Connective Tissue | Other | 25 | 7046 |
| Artefact | Other | 24 | 6862 |
| Fat | Other | 10 | 335 |
| Blood | Other | 10 | 1142 |
| Tumor Necrosis | Other | 7 | 798 |
| Tumor Stroma | Other | 6 | 555 |
| Scar | Other | 1 | 31 |

The model takes as input a 1536-dimensional feature vector per tile (extracted using H-optimus-1). Its architecture consists of a multi-layer perceptron (MLP) with three hidden layers of sizes 512, 256, and 128, using batch normalization, ReLU activation, and dropout (rate 0.3). It was trained for 50 epochs using the Adam optimizer, with a learning rate of 1e-4 and weight decay of 1e-5. To mitigate class imbalance, we applied a stratified sampling strategy during training, ensuring a more balanced representation of each class in every fold.

#### d. GeX-Pred Training: A Multiple Instance Learning (MIL) framework

Gex-Pred is a MIL model trained on the VESPER cohort to predict the expression of 848 subtype-associated genes in MIBC from histological tiles, using macro-dissection-level RNA-seq profiles as weak supervision. In the MIL framework, each macro-dissected region is treated as a bag of tiles (instances). Tile representations are aggregated into a slide-level embedding via a weighted sum, which is then used to predict bulk RNA expression. This strategy allows the model to leverage slide-level supervision while handling the absence of fine-grained annotations.

Formally, to predict bulk gene expression **y** from WSIs, each slide is first divided into *N* tiles, and each tile *i* is encoded into a feature vector **t**_*i*_ ∈ ℝ^*d*^. Tile embeddings are aggregated into a slide-level representation via an attention-based weighted sum:

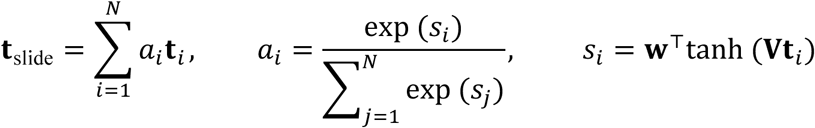

where **s**_*i*_ is the raw attention score for tile *i*, *a*_*i*_ is the normalized attention weight and **V** and *w* learnable parameters. The slide representation **t**_slide_ is then passed through a neural network *f*_o_) to predict the bulk RNA expression profile **y^**= ∈ ℝ*^G^*, with *G* = 848 genes:

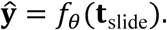

During training, 200 tiles were uniformly sampled from each region to improve computational efficiency, increase sampling diversity, and reduce memory usage. Tile representations were obtained using a pre-trained feature extractor (H-optimus-1), and the architecture is a MLP with two hidden layers of 512 neurons, each followed by batch normalization, ReLU activation, and dropout (rate 0.3). Training was conducted for 300 epochs using the Adam optimizer (learning rate = 1e-4, weight decay = 1e-5) and the Mean Squared Error (MSE) loss. Performances were assessed via 10-fold cross-validation.

At inference time, all tiles from each region were used. To generate pseudo-bulk predictions for macro-dissected regions or whole tumors, tile features were first aggregated, and the model then predicted the gene expression profile for the entire region, yielding a robust bulk-like estimate. When generating spatially resolved maps, each tile was individually assigned a predicted expression profile, enabling visualization of intratumoral gene expression and molecular subtype distribution.

For evaluation on external cohorts, predictions were obtained by ensembling the outputs of the 10 cross-validated models: the predicted gene expression was averaged across the 10 models, yielding a robust consensus prediction. When applying the model to whole-slide images (WSIs), MIBC-Detect was used beforehand to restrict inference to tiles predicted as MIBC. Based on the predicted expression profiles, a second model (MIBC-Clf) was then trained to assign consensus molecular subtypes at the tile level.

#### e. MIBC-Clf Training

We designed MIBC-Clf, a classification model that takes as input the gene expression profiles generated by GeX-Pred and outputs molecular subtype predictions. While a direct application of the consensus classifier would have been possible, MIBC-Clf was introduced to explicitly account for uncertainties inherent in the predicted expression profiles. In particular, while some genes are accurately predicted, others exhibit higher levels of uncertainty. By incorporating this additional modeling step, MIBC-Clf enhances robustness in downstream classification tasks.

MIBC-Clf is a multilayer perceptron (MLP) comprising three hidden layers with 512, 256, and 128 neurons, respectively. Each hidden layer is followed by batch normalization, ReLU activation, and dropout (dropout rate = 0.3). The model was trained for 200 epochs using the Adam optimizer (learning rate = 1×10⁻⁴, weight decay = 1×10⁻⁵) with Cross-Entropy (CE) loss. Performance was evaluated through 10-fold cross-validation. To enhance stability, the model was first pre-trained on synthetic data, where consensus classifier predictions were used as ground truth labels. Subsequently, the model was fine-tuned on GeX-Pred outputs, with GeX-Pred parameters frozen during this phase.

#### f. Gene-subtype association analysis for prediction performance

To associate each gene *i* with a consensus molecular subtype, we used the reference expression profiles (centroids) *m_c,i_* defining the consensus classifier subtypes *c* = 1, …, *C*. For each gene, we assigned it to the subtype showing maximal expression:

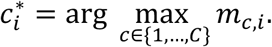

This gene-to-subtype assignment allowed us to assess Gex-Pred’s prediction performance at the gene level in relation to subtype-specific molecular programs.

### 3. Spatial Transcriptomics (ST)

#### a. Preprocessing of ST Data

For the comparison between predicted spatial gene expression and ST measurements, we resampled the ground truth resolution to match the prediction scale. The original ground truth data (VisiumHD) were acquired at 2 µm bins, which is much finer than the tile resolution (112 µm). To enable direct comparison, individual bins were aggregated into 100-µm patches by summing their expression values. Tumor patches were then identified by clustering patch-based expression profiles within each slide. To identify patches comprising tumor cells, clusters were scored using three predefined gene signatures (Basal, Luminal, and Epithelial) derived from CellMarker 2.0 [38]. If at least one of the signature scores exceeded a defined threshold, the cluster was retained. Expression values were subsequently log2+1 normalized prior to applying the consensus molecular classifier.

Two Visium HD slides were excluded from the analysis: one due to e abnormally low UMI counts (∼11 UMIs per 8 µm bin), and one due to severe image blurring that did not meet the minimal quality criteria required for morphology-based AI predictions.

#### b. Spatial Gene expression Prediction and consensus classification

To evaluate spatially resolved gene expression from histological predictions against ST, we applied our DL models directly to the whole-slide images corresponding to the VisiumHD data. For each 100-µm patch defined during ST preprocessing, a corresponding 112-µm image tile was extracted. Each tile was assigned both a predicted gene expression profile with Gex-Pred and a classification probability with MIBC-Clf. This approach enabled a direct, resolution-matched comparison between the spatial distributions predicted by Gex-Pred/MIBC-Clf and the expression-based classification measured by ST. Based on this alignment, we calculated Pearson correlations between predicted and measured gene expression, as well as ROC-AUC values to assess classification performance.

### 4. Model Evaluation and Benchmarking

#### a. Benchmark of foundation models for gene expression prediction

We benchmarked a series of foundation models using cross-validation on the VESPER cohort for transcriptomic prediction tasks. Supplementary Table 2 summarizes each model’s performance in predicting gene expression as well as molecular subtype classification. For simplicity of the benchmark, we focused the training exclusively on the gene expression prediction task (Gex-Pred). To obtain a measure of molecular subtype classification performance, we directly applied the consensus MIBC classifier to the predicted gene expression profiles. This approach allows us to fairly compare foundation models on transcriptomics prediction while assessing their downstream utility for molecular classification without retraining separate classifiers. All models were evaluated using the same downstream architecture for Gex-Pred and training pipeline to ensure a fair comparison (see Supplementary Section 2.d).

**Supplementary Table 2:** Benchmark of foundation models for transcriptomics prediction and molecular subtyping. Metric for transcriptomics prediction is median Pearson correlation coefficient (PCC) across genes and for molecular subtype classification accuracy ± standard deviation across cross-validation folds. All models were fine-tuned and evaluated using the same downstream architecture (MLP as described in Supplementary Section 2.d) and the same training pipeline to ensure a fair comparison. The table also reports training characteristics, including the number of WSIs used for pretraining and the corresponding resolution.

| Model | #WSIs for Pretraining + Resolution | Gene Expression (Median PCC) | Molecular Classif. (Accuracy) |
| --- | --- | --- | --- |
| H-optimus-1 | 1 000 000+ (20X) | **0.462** | **0.742** (+/- 0.05) |
| H-ptimus-0 | 500 000+ (20X) | 0.443 | 0.709 (+/- 0.08) |
| UNiv2 | 350 000+ (10-20X) | 0.449 | 0.708 (+/- 0.06) |
| Uni | 100 000 (10-20X) | 0.429 | 0.670 (+/- 0.06) |
| Phikon-v2 | 58 359 (20X) | 0.402 | 0.638 (+/- 0.07) |
| Phikon | 6 093 (20X) | 0.413 | 0.681 (+/- 0.05) |
| Ctranspath | 32 320 (10X) | 0.392 | 0.604 (+/- 0.07) |

The foundation models differ primarily by the number of whole-slide images (WSIs) used for pretraining. As expected, the more recent models trained on larger datasets show improved performance. For example, H-optimus-1, pretrained on over one million WSIs at 20X resolution, achieves the highest median gene expression PCC of 0.462 and the best molecular classification accuracy of 0.742 (±0.05). In contrast, earlier models like Ctranspath, pretrained on 32,320 WSIs at 10X resolution, perform notably worse with a median gene expression PCC of 0.392 and classification accuracy of 0.604 (±0.07). This clear trend highlights the importance of large-scale, high-resolution pretraining for foundation models in this domain.

Based on these results, we selected the top-performing foundation model, H-optimus-1, for all downstream analyses and results presented in this paper. Supplementary Figure 2 shows GeX-Pred predictions on the VESPER cohort under cross-validation, comparing predicted expression values with their corresponding targets. The results highlight a strong concordance for the top 20 genes, indicating that the model can reliably capture gene-specific expression patterns within the training cohort.

**Supplementary Figure 2:**
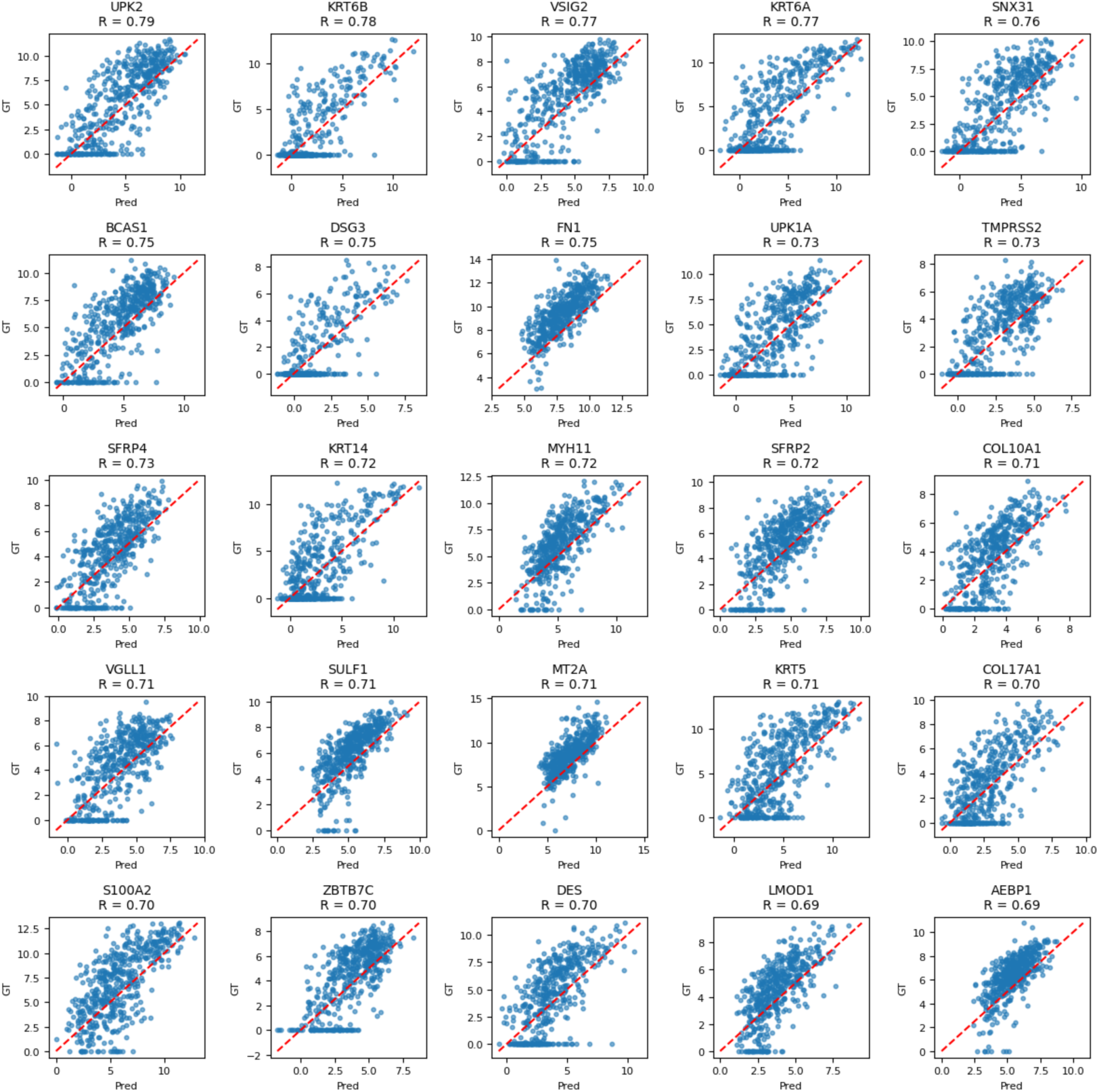
GeX-Pred on the Vesper cohort. Predicted versus target expression values for the 20 best-predicted genes under cross-validation, showing strong concordance within the training cohort.

#### b. Comparison of Direct Classification and Indirect Gene Expression Prediction for Molecular Subtyping

We compared two strategies for predicting molecular subtypes from histopathology whole-slide images (WSIs). In the first setup (*Direct Classifier*), the models directly predict the subtype labels from the WSIs. In the second setup (*Gex-Pred + MIBC-Clf*), the models first predict transcriptomic expression vectors, which are then used as input to a separate classifier for the final subtype assignment. Supplementary Figure 3 presents the comparison results of the two approaches across molecular subtypes for the VESPER cohort.

**Supplementary Figure 3:**
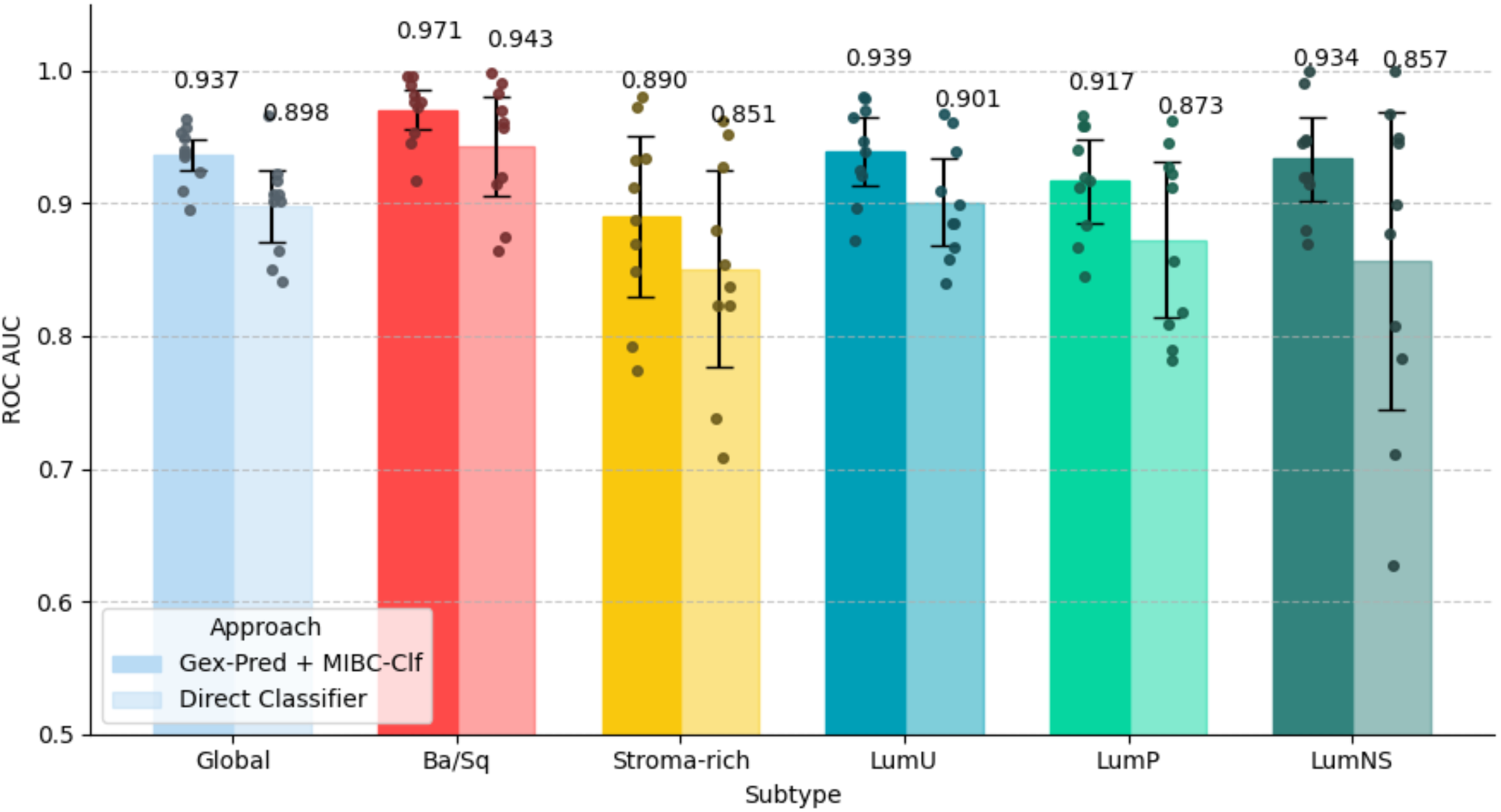
Comparison of direct classification and intermediate gene expression prediction for molecular subtyping from histopathology WSIs. Global and subtype-specific ROC AUC for the two prediction strategies obtained by cross-validation on the VESPER cohort.

Overall, the indirect approach leads to a clear improvement in classification performance, with the global test ROC AUC increasing from 0.898 ± 0.04 (Direct) to 0.937 ± 0.02 (GeX-Pred + MIBC-Clf). This gain is consistent across all molecular subtypes. The majority subtype Ba/Sq, which includes 180 cases, improves from 0.943 to 0.971 (a gain of 0.028), while the under-represented LumNS subtype, represented by only 32 cases, shows a larger improvement from 0.857 to 0.934 (a gain of 0.077). This difference highlights how the intermediate transcriptomic representation helps mitigate the impact of class imbalance.

The VESPER dataset is notably imbalanced: some subtypes like LumNS are represented by only 32 cases, making direct training challenging. By regressing gene expression profiles first, the model benefits from the continuous nature of transcriptomic data, providing a richer supervision signal and enabling multi-task learning. This strategy helps capture both transcriptomic and subtype-specific features more robustly, improving predictions especially for rare classes that otherwise suffer from limited examples and harder decision boundaries.

Based on this significant and consistent improvement, we retained the indirect approach (Gex-Pred + MIBC-Clf) for all experiments in this study.

#### c. Comparison of model performance on TURBT and Cystectomy specimens

Our models, GeX-Pred and MIBC-Clf, were trained on the VESPER cohort, which exclusively included TURBT specimens. Here, we aimed to demonstrate that our approach generalizes well to cystectomy specimens (treatment-naïve), while maintaining consistent performance. In the COBLANCE Frozen cohort, patients had either TURBT samples (n = 49) or cystectomy samples (n = 98, all NAC-naïve). The cohort was therefore divided into two groups to compare model performance between TURBT and cystectomy specimens.

As shown in Supplementary Figure 4, gene expression prediction achieved slightly higher accuracy for TURBT samples (PCC = 0.51) compared to cystectomy samples (PCC = 0.48). Conversely, molecular subtype classification yielded a ROC AUC of 0.801 for TURBT and 0.876 for cystectomy. These results demonstrate that our models generalize effectively to cystectomy samples.

**Supplementary Figure 4:**
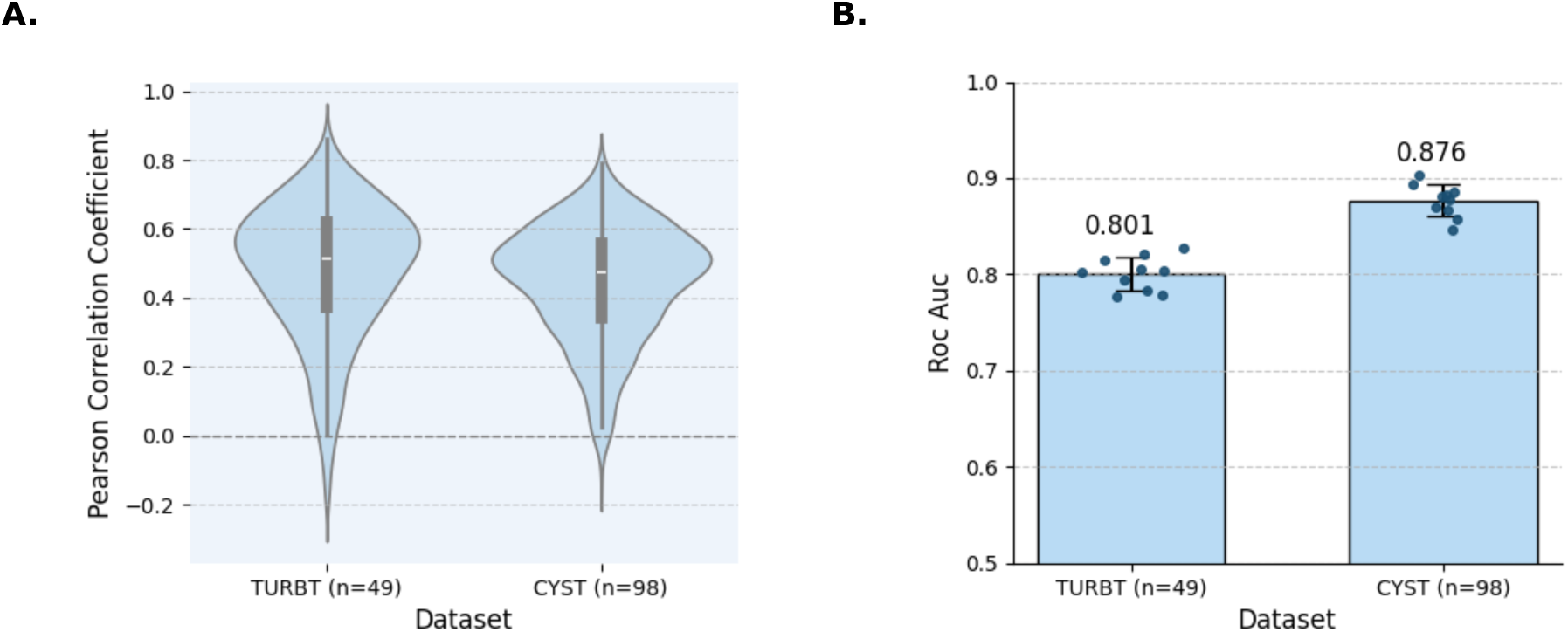
Comparison of model performance on TURBT and cystectomy (CYST) specimens in the COBLAnCE Frozen cohort. Patients were divided into two subgroups according to specimen type (TURBT or CYST). **A.** Gene expression prediction performance (PCC) between bulk RNA-seq and GeX-Pred. **B.** Molecular subtype classification performance (ROC AUC) between RNA-seq ground truth and MIBC-Clf predictions.

#### d. Additional ST prediction examples across genes and slides

To complement Figure 3, Supplementary Figure 5 presents additional visualizations of the ST predictions across all available slides and for a broader set of genes. These plots further highlight the consistency between predicted and measured spatial expression patterns for the top genes.

#### e. MIBC-Detect performances

MIBC-Detect achieved excellent performance in classifying histological tiles into three categories (non-tumoral, NMIBC, and MIBC). The model reached a ROC-AUC of 0.978, indicating an excellent separation between non-tumoral and tumoral tissues. Supplementary Figure 6 illustrates these results: panel A reports the overall classification metrics, while panel B shows representative prediction maps (a TURBT and a cystectomy). These maps highlight the model’s ability to effectively exclude large sets of non-tumoral tiles, thereby focusing the detection on clinically relevant MIBC regions.

#### f. Interpretability and high score tiles

Understanding how the model captures and distinguishes histological features across molecular classes is essential for interpretability. To investigate which morphologies the model associated with each class, we examined the VESPER cohort and selected the 20 tiles with the highest scores for each subtype. To ensure diversity, only the top-scoring tile from each patient was retained. Representative examples are shown in Supplementary Figure 7A. Some morphological patterns highlighted by the model were consistent with those previously described in the literature. Specifically, top scoring Ba/Sq tumor tiles often showed squamous differentiation, characterized by the presence of large polygonal cells with intercellular bridges and/or keratinization [7,12]. Others displayed cells with prominent nucleoli, as previously reported [17]. In addition, inflammatory cells were often observed within the tumor microenvironment [7]. Many top LumU tiles harbored micropapillary features, as previously suggested [12].The Stroma-rich subtype tiles presented abundant fibroblastic or smooth muscle connective tissue, in which tumor cells appear more scattered [7]. Finally, many LumP tumor tiles focused on areas of papillary architecture, composed of thin fibrovascular cores covered by urothelial carcinoma cells [7].

**Supplementary Figure 5:**
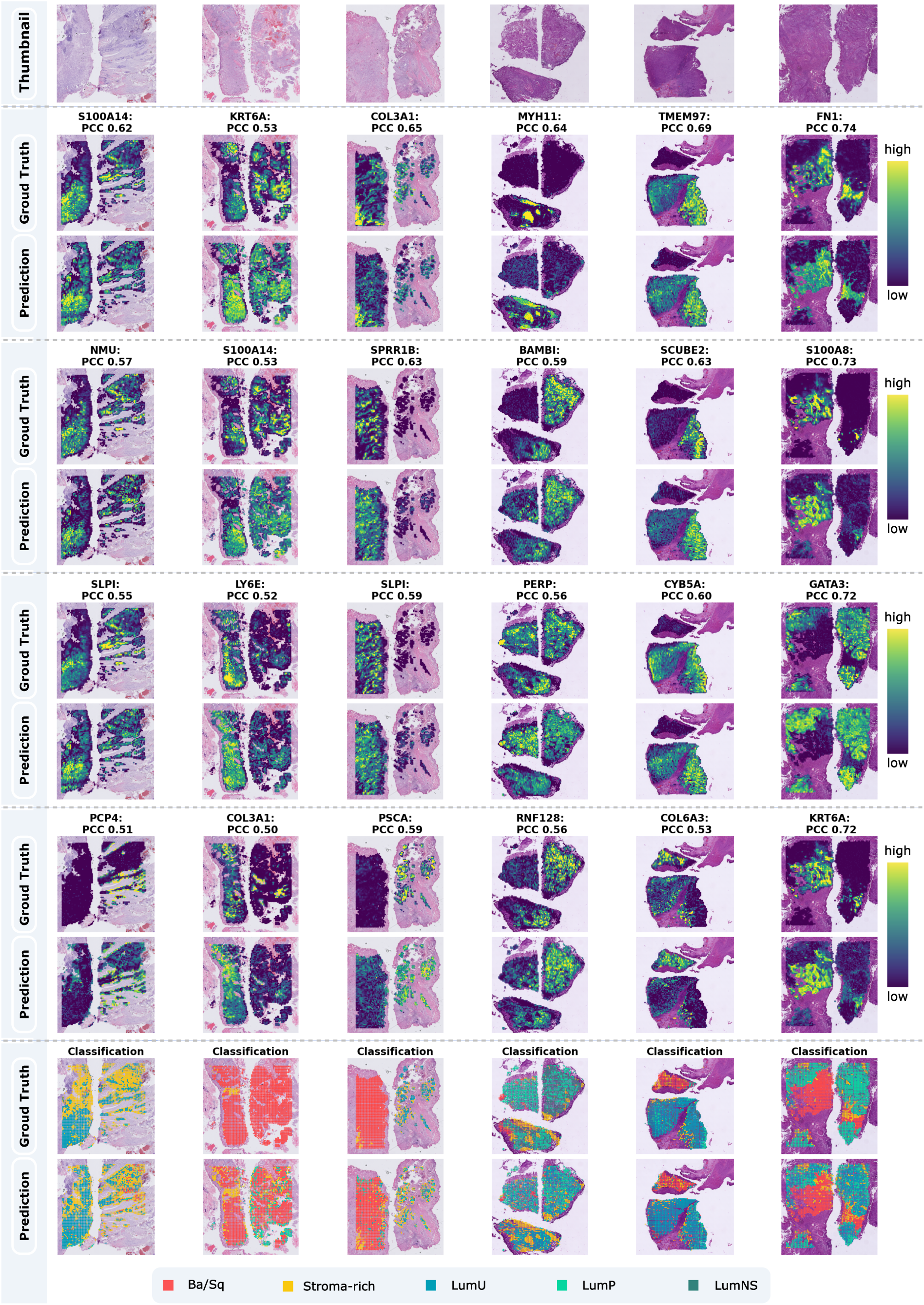
Supplementary visualizations of spatial transcriptomics predictions across all six Visium slides. The first row shows the thumbnails of all Visium slides used in this analysis. For each slide, four representative genes were selected based on a trade-off between prediction performance and diversity (i.e., low correlation between genes). For each gene, the first row corresponds to the measured spatial transcriptomic signal (*GT*), and the second row to the predicted expression map (*Prediction*). At the bottom, the corresponding molecular classification maps are displayed for each slide, with the first row showing the measured (*GT*) and the second row the predicted (*Prediction*) molecular subtype distributions.

**Supplementary Figure 6:**
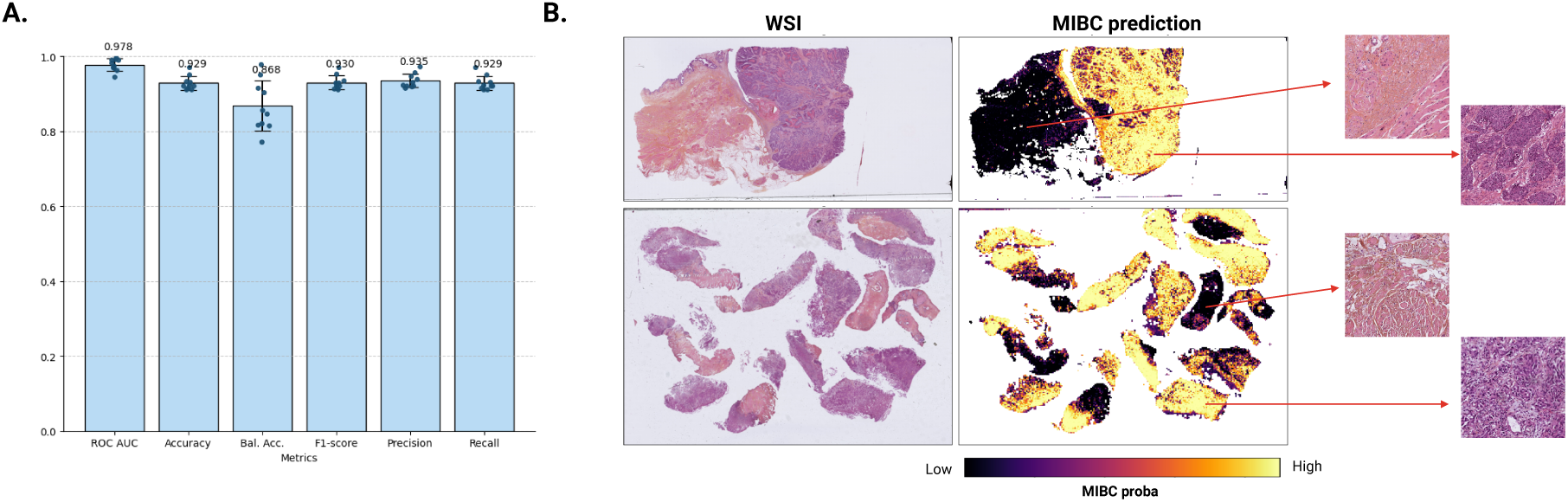
Performance of MIBC-Detect. **A.** Classification metrics obtained through cross-validation on the Vesper annotations. **B.** Prediction maps generated by MIBC-Detect on whole-slide images, including MIBC probability maps and representative tiles predicted as MIBC or non-MIBC.

#### g. Robustness to staining variation

The proposed DL-based approach aims to identify molecular subtypes directly from histology, with the clinical objective of prognostic stratification. To demonstrate its potential for clinical implementation, it is crucial to establish that the model is robust across institutions and compatible with variations in staining, scanners, and protocols. We therefore assessed prediction consistency under staining variation using the COBLANCE cohort, which includes 142 cases initially stained in 13 different centers and subsequently restained at Institut Curie on consecutive slides. Prediction consistency was assessed by computing, for each case, the proportion of tiles assigned to each subtype and calculating the correlation between predictions on the original slide and the restained slide. Model predictions were highly consistent, with a 95.4% median-correlation between original and Curie-stained slides. A subset of slides, along with their predictions across both stainings, is presented in Supplementary Figure 7B.

**Supplementary Figure 7:**
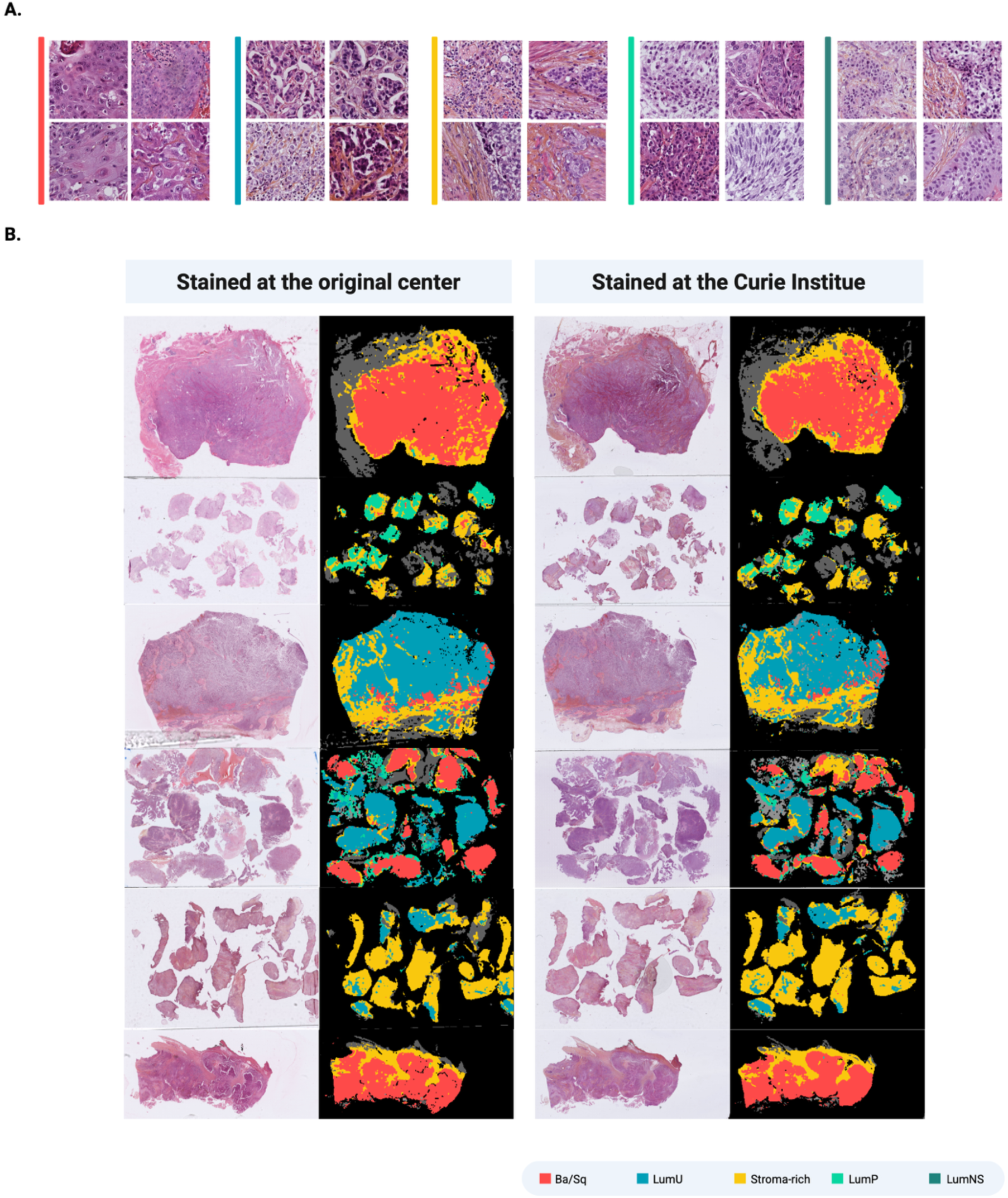
Robustness to staining variation and representative high-score tiles. **A.** Robustness of molecular subtyping across staining protocols. Representative examples showing consistent spatial prediction maps between original slides stained in 13 external centers and a consecutive slide restained at Institut Curie. **B.** Representative tiles with the highest scores for each molecular subtype of the consensus classification in the VESPER cohort. One top-scoring tile per patient was retained to ensure diversity.

#### h. Survival analysis of macro-dissections stratified by Ba/Sq versus non-Ba/Sq

Predictions from our model indicated that 41% of patients in the VESPER cohort harbor a Ba/Sq component. Kaplan–Meier survival analysis revealed that tumors predicted as Ba/Sq, whether pure or mixed, were significantly associated with poorer progression-free survival compared to tumors lacking a Ba/Sq component (log-rank p PFS = 0.015, OS = 0.016) (Supplementary Figure 8A,B). This finding is consistent with RNA-seq–based stratification (log-rank p PFS = 0.001).

**Supplementary Figure 8:**
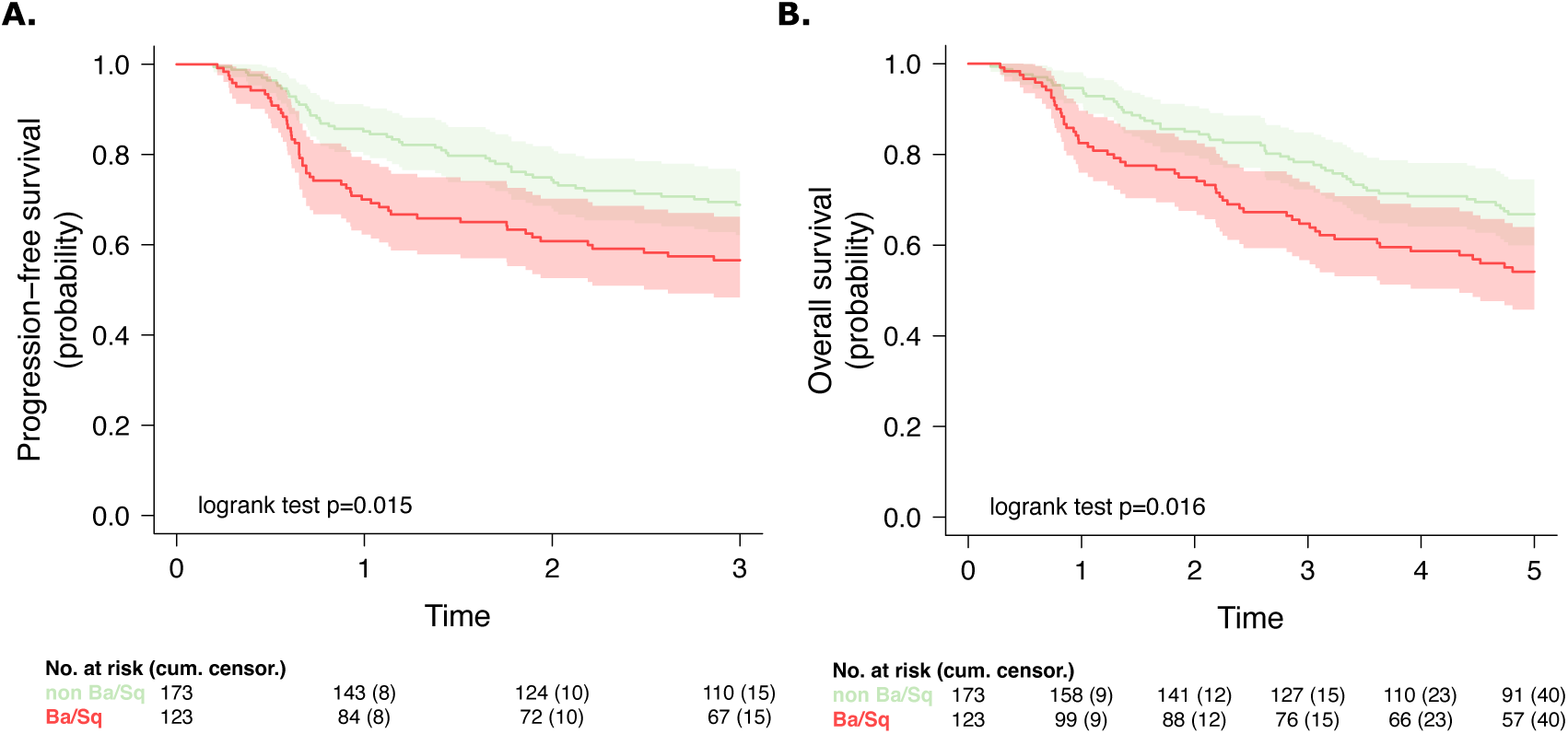
Survival analysis of Ba/Sq versus non-Ba/Sq tumors in the VESPER cohort based on predictions from macro-dissected regions. Kaplan–Meier curves for progression-free interval (**A**, PFI) and overall survival (**B**, OS) based on deep learning predictions from histology.

